# Inhibitory and Transport Mechanisms of the Human Cation-Chloride Cotransport KCC1

**DOI:** 10.1101/2020.07.26.221770

**Authors:** Yongxiang Zhao, Jiemin Shen, Qinzhe Wang, Ming Zhou, Erhu Cao

## Abstract

Secondary active cation-chloride cotransporters (CCCs) catalyze electroneutral symport of Cl^−^ with Na^+^ and/or K^+^ across membranes^1,2^. CCCs are fundamental in cell volume homeostasis, transepithelia ion movement, maintenance of intracellular Cl^−^ concentration, and inhibitory synaptic transmission^3–6^. K^+^-Cl^−^ cotransport 1 (KCC1) was first characterized in red blood cells and later in many other cell types as a crucial player in regulatory volume decrease in defense against cell swelling upon hypotonic challenges^7,8^. Here we present two cryo-EM structures of human KCC1: one captured in an inward-open state and another arrested in an outward-open state by a small molecule inhibitor. KCC1 can surprisingly adopt two distinct dimeric architectures via homotypic association of different protein domains and conversion between these two forms of dimers may entail dynamic formation and rupture of two interdigitating regulatory cytoplasmic domains. The inhibitor wedges into and forces open an extracellular ion permeation path and arrests KCC1 in an outward-open conformation. Concomitantly, the outward-open conformation involves inward movement of the transmembrane helix 8 and occlusion of the intracellular exit by a conserved short helix within the intracellular loop 1. Our structures provide a blueprint for understanding the mechanisms of CCC transporters and their inhibition by small molecule compounds.

## Introduction

Secondary active cation-chloride cotransporters (CCCs) harness the Na^+^ and/or K^+^ gradients created by Na^+^-K^+^-ATPase pumps to move Cl^−^ into or out of cells in an electroneutral fashion^1,2^. CCCs can be divided into two clades: three Na^+^-dependent Na^+^-(K^+^)-Cl^−^ (NCC and NKCC1-2) and four Na^+^-independent K^+^-Cl^−^ (KCC1-4) cotransporters. CCCs play fundamental roles in a multitude of biological processes such as transepithelial ion movements, cell volume homeostasis, regulation of intracellular Cl^−^ concentration ([Cl^−^]i) and neuronal excitability^3–6^. A K^+^-Cl^−^ cotransport mechanism was initially characterized in red blood cells as a crucial effector of regulatory volume decrease upon cell swelling^7,8^. In red blood cells and later discovered in many other cell types, in response to hypotonic challenge, KCCs extrude Cl^−^ and K^+^ to defend against cell swelling via concomitant obligatory water efflux. Indeed, inhibiting KCCs activity was shown to prevent sickle cell dehydration and thus provides a promising therapeutic strategy to treat sickle cell anemia^9,10^. In neurons, KCC2 functions as a major Cl^−^ extruder to maintain [Cl^−^]i below electrochemical equilibrium such that inhibitory neurotransmitters (e.g., γ-aminobutyric acid; GABA) stimulate Cl^−^ influx via pentameric ligand-gated Cl^−^ channels and lead to hyperpolarization of neurons^11^. Importantly, mutations in KCC2 and KCC3 cause brain disorders such as epilepsy, seizures, and sensorimotor neuropathy with agenesis of corpus callosum^12–19^. Given the pivotal roles of KCCs in inhibitory synaptic transmission, KCCs are emerging attractive targets for developing therapeutic strategies to restore GABA inhibition for the treatment of various brain disorders and psychiatric conditions^11,20–23^.

Several cryo-EM structures of CCC transporters have been recently reported and provided valuable insights into their architectural design principles and mechanisms of ion binding^24–27^. These structures demonstrate that the CCC transport core consists of two inverted repeats of five-helices bundle (i.e., TM1-TM5 and TM6-TM10), with the remaining two helices (TM11 and TM12) optionally contributing to dimeric assembly in addition to cytoplasmic C-terminal domain or extracellular domain. The TM1 and TM6 helices lie at the center of ion transport path and break **α**-helical geometry roughly at the middle of lipid bilayer where ion binding sites are organized around these discontinuous hinge regions. Unfortunately, all CCC structures reported to date were captured in an inward-open state and cannot to be simply extrapolated to understand other transport states (e.g., outward-open and occlude states) that a CCC transporter must sample to shuttle ions across membrane. Moreover, several members of the CCC family are inhibited by loop and thiazide diuretic drugs for the treatment of hypertension and edema or by tool compounds developed for probing their structures and functions^28–31^. We are yet to determine a structure of a CCC transporter bound with such an inhibitor which would reveal their sites and mechanisms of action and facilitate medicinal chemistry and computational docking efforts to further improve specificity and potency of these drugs.

Here we reported two cryo-EM structures of human KCC1 that represent the transporter trapped in an inward-open state and arrested in an outward-open state by the VU0463271 inhibitor, respectively. We showed that KCC1 can adopt two drastically distinct dimeric architectures by assembling through different structural domains, raising the possibility that posttranslational modification such as phosphorylation or binding of cellular factors may preferentially regulate formation of these two types of dimers, and consequently, transport activity. We further showed that the inhibitor wedges into the extracellular ion transport path, breaking ionic gating interactions at the extracellular entryway and leading to opening of an extracellular vestibule. Concomitantly, inward movements of TM8, coupled with a concerted displacement of a highly conserved short helix within the first intracellular loop, occlude the cytoplasmic exit as seen in the inward-open KCC1 structure. Our structures shed lights on the conformational landscape of KCC1 as it proceeds along a transport cycle and may facilitate rational development of small molecules to modulate KCCs activity for the treatment of various brain disorders.

## Results

### Human KCC1 adopts two distinct dimeric architecture

We have determined two structures of the full-length human KCC1 transporter by single-particle cryo-EM: one in 150 mM KCl and Lauryl Maltose Neopentyl Glycol (MNG-3) detergent at 3.25 Å resolution and another in 150 mM NaCl and MNG-3 detergent further supplemented with the VU0463271 inhibitor (referred to VU hereafter) at 3.63 Å resolution (**Figure 1 and Extended Data Figure 1-6**). In keeping with previously reported CCC structures^24–26^, the human KCC1 transporter also assembles as a dimer in both conditions, albeit assuming drastically distinct overall architecture due to association via different inter-subunit interfaces (**Figure 1 and Extended Data Figure 7**; also see below). In both structures, each KCC1 subunit consists of 12 transmembrane helices (TM1-TM12), in which TM1-TM5 and TM6-TM10 are two inversely oriented helix bundles related by a pseudo two-fold symmetry axis parallel to the membrane plane. Moreover, in both KCC1 structures, TM1 and TM6 helices split into two half helices (TM1a and TM1b; TM6a and TM6b) as they become discontinuous with a broken **α**-helical geometry roughly half way across the lipid bilayer.

**Figure 1.**
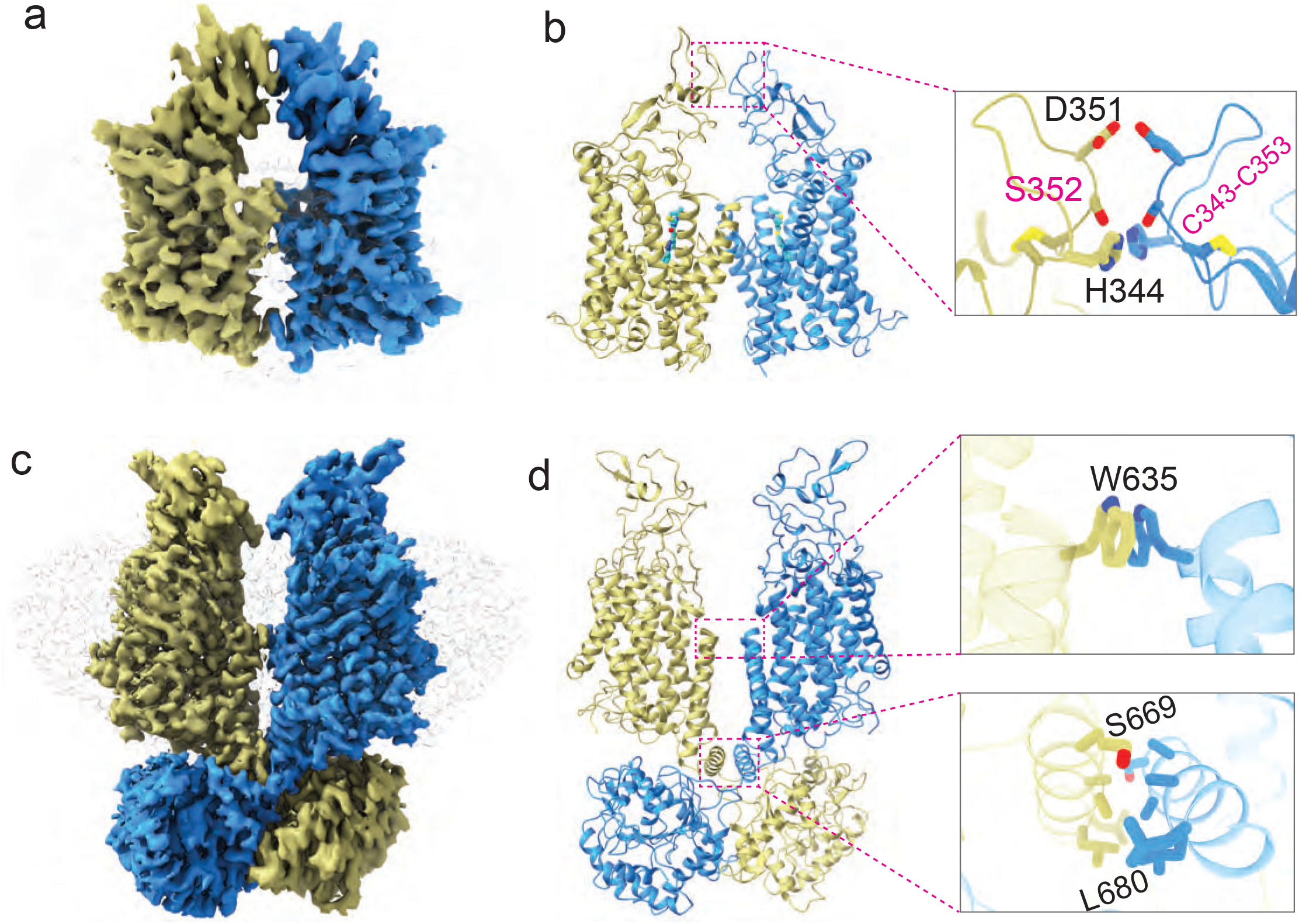
Human KCC1 adopts two dimeric architecture. (a) Side view of human KCC1 bound with the VU0463271 inhibitor. Individual transporter subunits are color-coded. Densities of detergent micelle (grey) are rendered semi-transparent. (b) Ribbon representation of KCC1 dimer bound with the inhibitor is shown in the same orientation as in (a) with an enlarged view highlighting the dimer interface between the large extracellular domains. (c) Side view of human KCC1 determined in the presence of 150 mM KCl. Individual transporter subunits are color-coded. Densities of detergent micelle (grey) are rendered semi-transparent. (d) Ribbon representation of KCC1 dimer in the presence of 150 mM KCl is shown in the same orientation as in (c) with enlarged views of two dimer interfaces.

Despite that these two KCC1 dimers share overall structural similarity on a single subunit level when cytosolic domains are excluded (**Figure 2a**), two KCC1 subunits can surprisingly assemble into two drastically distinct dimeric architectures (**Figure 1 and Extended Data Figure 7d**). In the 150 mM KCl (or Apo) condition, the KCC1 dimer resembles structures of zebrafish NKCC1 and KCC3 (PDB code: 6Y5R) where an inverted V-shaped helix-turn-helix structure formed by TM11 and TM12 helices constitutes the principal dimeric interface within the lipid bilayer in addition to the interdigitating cytosolic C-terminal domains. In particular, an **α**-helix (^666^Arg-Glu^682^), which immediately follows TM12, runs almost parallel to the inner membrane and establishes extensive hydrophobic interactions with the same helix oriented in an opposite direction from a second KCC1 subunit. In the VU-bound KCC1 structure, however, cytosolic domains are not resolved in our map, consistent with reference free 2-dimensional (2D) class averages that also only show a cloud of fuzzy densities for the cytosolic domains (**Extended Data Figure 1**). We suspect that cytosolic domains may not participate in dimer assembly in the VU-bound KCC1 structure. Instead, a large extracellular domain (ECD), formed by a stretch of ~ 120 residues between TM5 and TM6, participates in homotypic interactions with the same structure from a second KCC1 subunit. In particular, residues (i.e., His344, Asp351, Ser352) protrude from a finger-like loop, a structure that is possibly rigidified by a conserved disulfide bond found in all KCC isoforms, pointing their side chains toward the central two-fold axis to establish polar contacts with the same set of residues from another subunit. Intriguingly, no direct protein contacts exist between the two KCC1 subunits within the lipid bilayer in the VU-bound KCC1 structure. However, non-protein densities, which possibly represent co-purified endogenous lipids (or detergents), are found in the large void between the two subunits, suggesting that lipid-mediated association within the lipid bilayer may additionally stabilize KCC1 dimer within native membrane in addition to interactions among the large ECD. We noted that our VU-bound KCC1 adopts a similar dimeric architecture as previously reported KCC1 structures^26^, although relative orientation of two subunits, and consequently, inter-subunit interfaces are not identical in these two structures (**Extended Data Figure 7d**).

**Figure 2.**
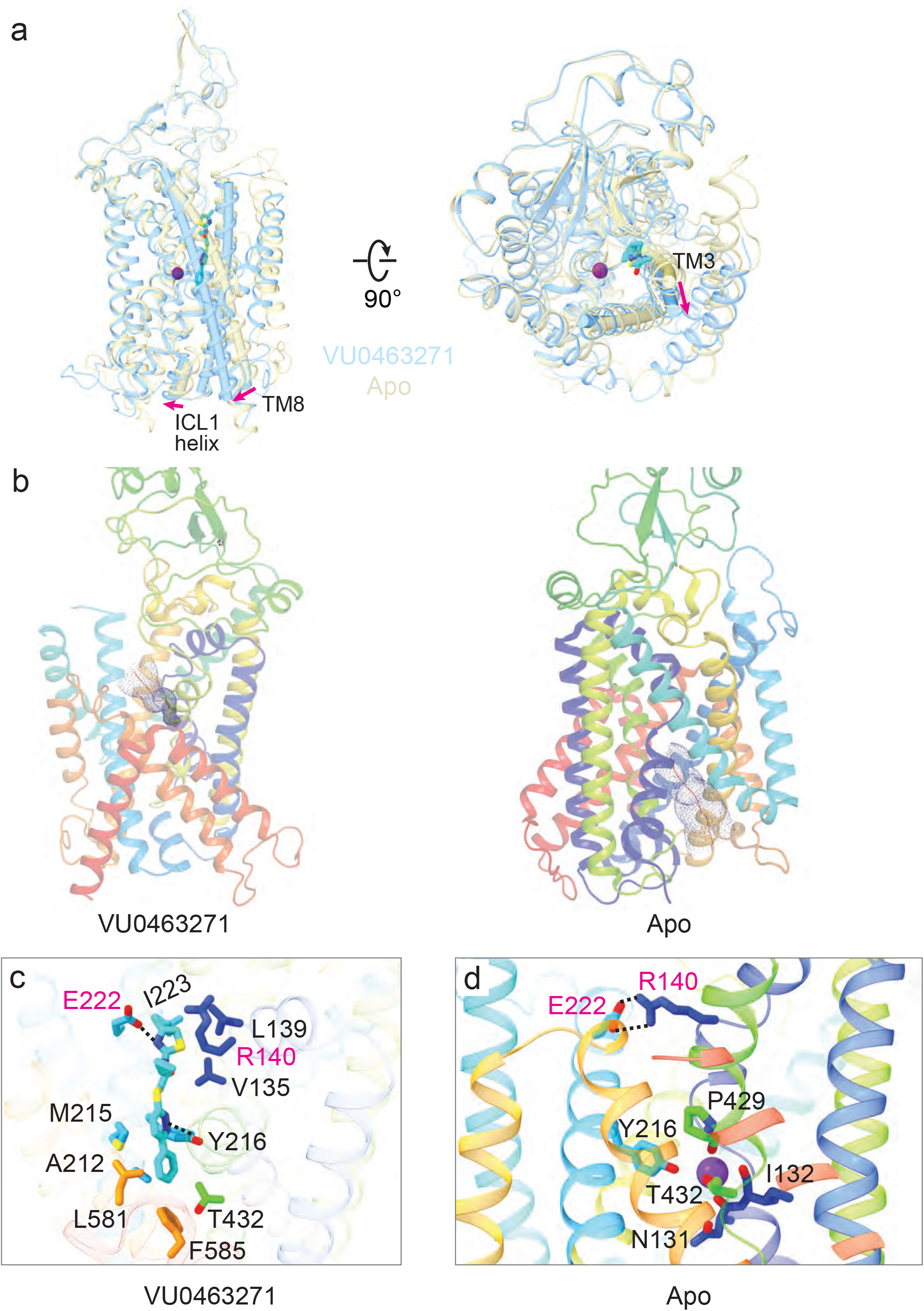
The VU0463271 inhibitor arrests KCC1 in an outward-open state. (a) Superimposition of KCC1 structures in VU0463271-bound and 150 mM KCl (apo) states. K^+^ ion and the inhibitor are shown as a purple sphere and sticks, respectively. Movements of TM3, TM8, and the ICL1 helix are highlighted with red arrows. (b) Solvent assessible pathways are mapped with the HOLE program, highlighting an extracellular vestibule in the VU0463271-bound KCC1 structure if the inhibitor is removed and an intracellular exit in the apo KCC1 structure. (c) The VU0463271 breaks the ionic extracellular gate formed between E222 and R140 and interacts with residues along the extracellular ion permeation pathway. (d) The extracellular gate and K^+^ binding site are intact in the KCC1 structure determined without the inhibitor (apo) state.

It is quite intriguing that human KCC1 can adopt two drastically distinct dimeric architecture that seems to depend on whether the VU inhibitor is present or not. When we docked the full-length structures of KCC1 (in 150 mM KCl) and KCC3 into the VU-bound KCC1 map, we found that the ECD and transmembrane domain roughly fit into the map without notable clashes between two subunits, but the two interdigitating C-terminal domains are now completely separated from each other (**Extended Data Figure 7**). Indeed, the shortest distance between two C-terminal domains would be approximately 25 Å in the VU-bound KCC1 dimer, assuming that the monomeric architecture is preserved in both conditions. This provides a plausible explanation why we could not resolve the cytosolic domain in the VU-bound KCC1 map as formation of this type of dimer possibly entails rupture of the extensive cytosolic interface as observed in KCC1 and KCC3 structures determined without any inhibitor. It remains to be determined whether these two types of dimers exist on native cells, but it has been shown that activation of NKCC1 upon phosphorylation leads to separation of two C-terminal domains based on Förster resonance energy transfer (FRET) measurements^32^, lending support to potential rupture of the extensive cytoplasmic dimer interface during the process of ion transport or phosphoregulation. Of note, the large ECD represents the most divergent structure in primary sequence among four KCC isoforms (**Extended Data Figure 8**), so it is unclear whether this domain can serve as a general dimerization module within all KCC transporters. Notwithstanding this important caveat, we found that KCC3 and KCC4 might be able to associate via the ECD domain as a set of analogous hydrophilic residues can similarly engage in homotypic interactions as seen in VU-bound KCC1 structure (**Extended Data Figure 7**). As the VU compound also inhibits transport activity of KCC3, future studies will determine whether it can also promote assembly of a second form of dimeric architecture resembling the VU-bound KCC1 structure. Taken together, our structural data showed that KCC1 exhibits remarkable plasticity in modes of dimeric assembly and future studies will determine whether conversion between these two dimeric forms is intimately associated with activation of CCC transporters by posttranslational modifications such as dephosphorylation or by engagement of cellular factors such as creatine kinases^33–35^.

### The VU0463271 inhibitor opens the extracellular gate

Several recently reported CCC structures of NKCC1, KCC1, KCC4, and KCC3 (PDB code: 6Y5R) have greatly advanced our mechanistic understanding of this physiologically and clinically important family of transporters^24–26^. Unfortunately, all these structures are captured in an inward-open conformation regardless of ionic conditions, thus transport-associated conformation changes and structure-function relationship remains incompletely understood without CCC structures determined in outward-open and occluded states. To fill this fundamental gap of knowledge, we sought to trap human KCC1 in a distinct state with the VU inhibitor as it competes with K^+^ for the same or overlapping binding site along the ion transport pathway of KCCs^31^, and also because the VU inhibitor has much larger size than the permeating K^+^ ion, it could squeeze into and physically open the extracellular entryway. Indeed, our VU-bound KCC1 structure showed that binding of the VU compound leads to opening of a large extracellular vestibule that would allow for unobstructed access of K^+^ and Cl^−^ ions to the central ion binding sites organized around the discontinuous hinge regions of TM1 and TM6 helices (**Figure 2**). In contrast, the intracellular exit is occluded (also discussed below), demonstrating that the VU-bound map represents KCC1 in an outward-open state. On the other hand, KCC1 assumes an inward-open conformation in 150 mM KCl (or Apo) condition. Comparison of these two KCC1 structures showed that opening of the extracellular entryway involves outward displacement of TM3, TM4, TM8, TM9 and TM10 helices. These rigid body movements of helices were possibly triggered by direct engagement of the VU compound with residues on TM3 and TM10 that could conceivably exert force on and push these two helices away from the center of the extracellular vestibule, and consequently, cause a concerted outward displacement of the three remaining adjacent helices as well. Consistent with our structural findings, in human NKCC1, TM10 was also inferred to undergo significant movements during ion transport as several residues on TM10, when substituted to cysteine, exhibit state-dependent solvent accessibility and crosslinking^36^.

The VU0463271 compound, *N*-cyclopropyl-*N*-(4-methyl-2-thiazolyl)-2-[(6-phenyl-3-pyridazinyl)thio] acetamide, snugly fits into an extracellular pocket formed by the TM1b, TM6a, TM3, and TM10 helices, not only physically plugging the extracellular entryway, but also possibly preventing KCC1 from isomerizing into other states via steric hindrance (**Figure 2**). The VU compound establishes a multitude of polar and hydrophobic contacts with residues along the extracellular ion transport pathway. For instance, at the mouth of the extracellular entryway, the 4-methyl-2-thiazolyl group of VU0463271 wedges between Arg140 and Glu222 residues, essentially breaking the salt bridge interactions that are indispensable for closure of the extracellular gate as seen in the KCC1 structure determined in an inward-open state (**Figure 2**). Here, the 4-methyl-2-thiazolyl group interacts with residues Glu222 and Ile223 on TM3, as well as residues Val135 and Leu139 on TM1b via hydrogen bonding and hydrophobic packing interactions. Our structure also showed that the 4-methy group established hydrophobic interactions with Ile223, rationalizing previous structure-activity relationship studies which indicate that removal of this 4-methyl group diminishes potency of the compound. An analogous salt bridge pair exists at the extracellular gate of NKCC1^24,25^, suggesting that opening of the extracellular entryway may also entail disruption of these salt bridge interactions in NKCC1 as well. Deeper toward the central ion binding site, the phenyl-3-pyridazinyl group engages with a number of residues located on TM6a, TM3, and TM10 helices, including Tyr216 that would otherwise participate in coordination of a permeating K^+^ ion (**Figure 2 and Extended Data Figure 9**). Our VU-bound KCC1 structure showed that nitrogen atoms of the phenyl-3-pyridazinyl group may form hydrogen bonds with Tyr216, thus providing a plausible explanation why the potency of an inhibitory VU compound decreases with increasing concentration of extracellular K^+^ ion^31^.

In order to confirm the VU-binding site observed in our structure, we reconstituted wild type KCC1 and M215A mutant into liposomes to measure their sensitivity to the VU inhibitor using a thallium (a congener of K^+^) ion flux assay. We found that the M215A mutant exhibits diminished sensitivity to the VU inhibitor by ~ 65 folds, with an IC_50_ of 4.53 μM as compared to 69.8 nM determined for the wild type KCC1 (**Figure 3**). Of note, M382 in NKCC1, which is equivalent to M215 in KCC1, is essential for inhibition by loop diuretic bumetanide based on scanning mutagenesis studies^37^. Future structural studies will determine whether clinically prescribed loop diuretics also occupy an analogous packet in NKCC1 and NKCC2 transporters to block ion absorption in the ascending loop of Henle of nephrons in the kidneys. In other APC transporters, the extracellular entryway is also frequently targeted by many competitive or allosteric inhibitors, including tryptophan and anti-depressant citalopram that inhibit LeuT and serotonin transporter, respectively^38,39^.

**Figure 3.**
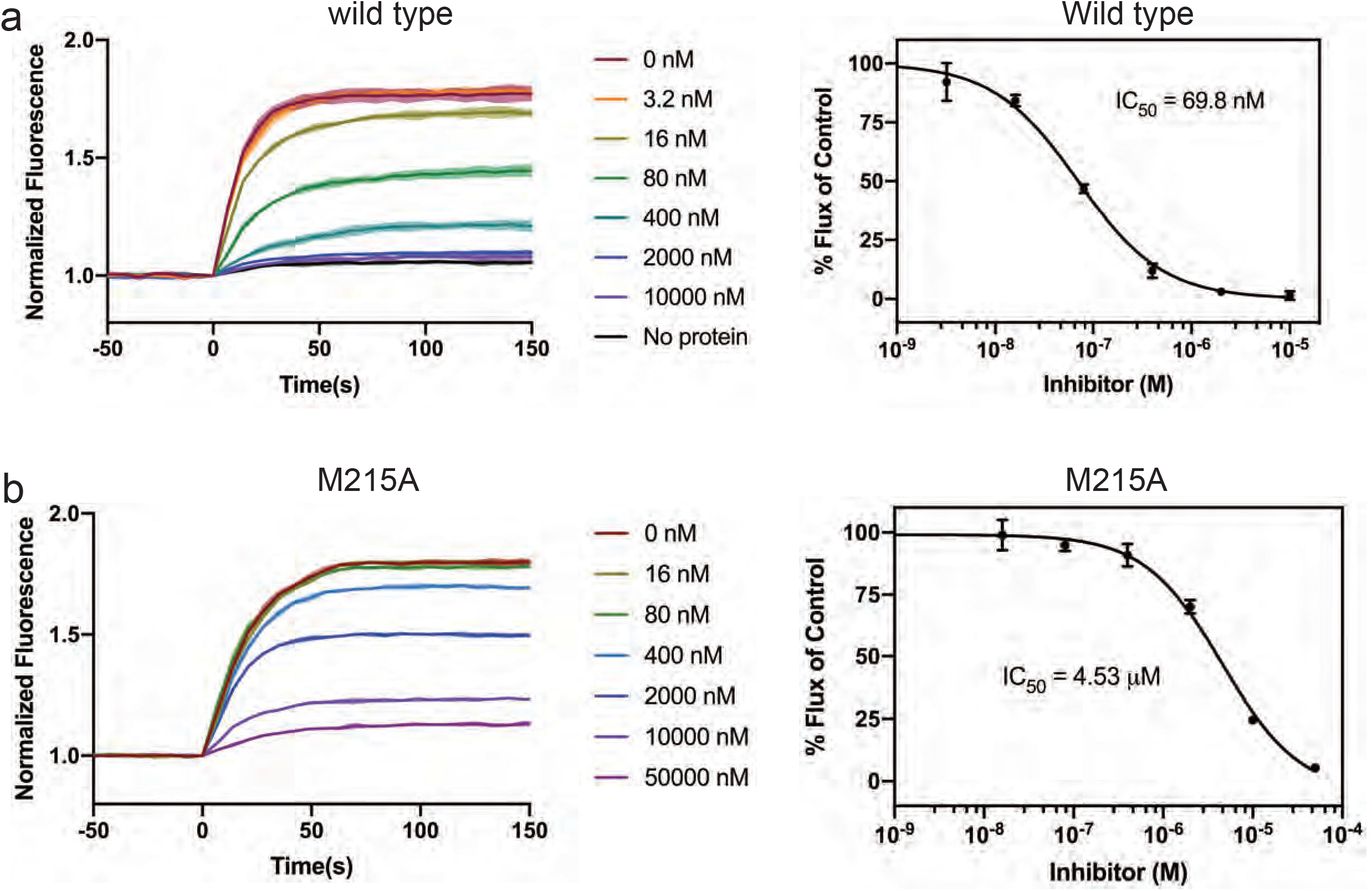
Mutation in the VU0463271 binding site affects inhibitory potency. (a) The wild type KCC1 is inhibited by VU0463271 with an IC_50_ of 69.8 nM measured in liposomes using a thallium ion flux assay. (b) KCC1 M215A mutant exhibited reduced sensitivity to the inhibitor with an IC_50_ of 4.53 μM.

### Intracellular gate

The classic alternate-access model dictates that substrate-binding site of a transporter is alternately exposed to opposite sides of the membrane as the transporter proceeds along a transport cycle^40^. Opening of an extracellular vestibule observed in our VU-bound KCC1 structure thus necessitates concomitant closure of the cytoplasmic exit. Indeed, comparison with the inward-open human KCC1 structure determined without the VU compound showed that a concerted inward movements of TM8 helix, the intracellular loop between TM6b and TM7, and the short helix within the strictly conserved first intracellular loop (ICL1) occlude the cytoplasmic exit (**Figure 2**). The most pronounced movements occur in TM8 and the short helix in ICL1, which are displaced by ~ 6.5 Å and 3.2 Å, respectively, as KCC1 isomerizes from an inward-open to an outward-open state. Such movements foster the establishment of new gating interactions that may contribute to closure of the cytoplasmic exit. For instance, Arg440 residing at the cytoplasmic end of TM6b participates in hydrogen bonding interactions with Gln521 and Thr524 located on TM8 in the VU-bound KCC1 structure **(Figure 4)**. In contrast, in the inward-open state, Arg440 is separated from Gln521 and Thr524 by as much as 10 Å measured as the shortest distance between their side chains. Of note, although these three residues are strictly conserved in all KCC isoforms and thus are likely crucial for isomerization of these transporters among distinct transport states, they are replaced by hydrophobic residues in NKCC1 (**Extended Data Figure 8**), suggesting that NKCC1, possibly other Na^+^-dependent clade of CCC transporters as well, may use a different set of residues to close intracellular exit.

**Figure 4.**
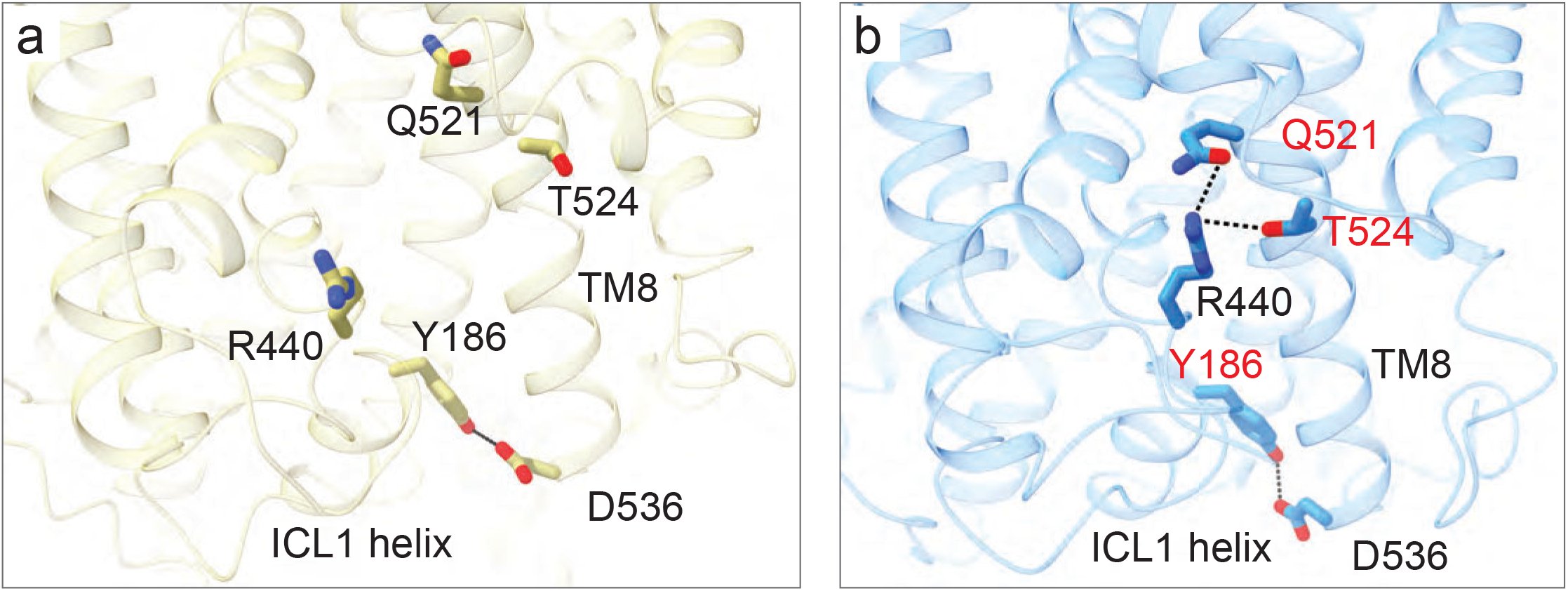
An intracellular gate in KCC1. (a) In the apo state, the intracellular gate is broken as R440 on TM6b is ~ 10 Å away from Q521 and T524 on TM8. (b) In the VU0463271-bound KCC1 structure, R440 establishes gating interactions with Q521 and T524 due to an inward movement of TM8. In both structures, the ICL1 helix is coupled to TM8 via hydrogen bonding interactions between Y186 and D536.

The ICL1 represents one of the most conserved region among of all CCC transporters, and in the case of NKCC transporters, missense mutations within this region have been shown to render transporters inactive, and consequently, development of Bartter’s syndrome in human^41,42^. In the VU-bound KCC1 structure, the ICL1 lies almost parallel to the inner leaflet of bilayer, intercalating among the cytoplasmic ends of TM2, TM3, TM6b, TM8, and TM10 and thus effectively plugging the intracellular vestibule. In particular, the short ICL1 helix appears to be directly coupled to TM8 via hydrogen bonding interactions between Tyr186 and Asp536, likely accounting for a concerted drastic displacement of both helices that leads to closure of the intracellular exit (**Figure 4**). Notably, opening of the intracellular exit in KCC1 does not involve hinge-bending motion of the TM1a helix as commonly observed in LeuT and neurotransmitter re-uptake transporters^43,44^.

## Discussion

Here we reported two cryo-EM structures of human KCC1: one captured in an inward-open state and another locked in an outward-open state by an inhibitor. Although a KCC1 structure trapped in an occluded state still awaits future structural and molecular modeling studies, we can begin to understand conformational changes and dynamic gating interactions that underlie ion transport by KCC1 and possibly by other CCC transporters as well. One unexpected finding emerged from our structural studies is that KCC1 isomerizes from outward-open to inward-open state without evoking drastic flexing motions of TM1 and TM6 half helices around the discontinuous hinge regions as commonly observed in LeuT and many other APC transporters^40,43–45^. Such subtle conformational changes associated with ion transport by CCC transporters possibly explain why they can catalyze much higher rates of substrate movement across membrane than many other APC transporters to which CCC transporters belong as previously noted^37^. Instead, we found that outward movements of TM3 pivoted at its cytoplasmic end, together with a slight displacement of TM10, result in opening of an extracellular pathway leading to the central ion binding sites (**Figure 5**). Although such displacements of helices were induced by the binding of VU inhibitor in our current structure, they may be similarly triggered by extracellular K^+^ and Cl^−^ in native cells as ions could conceivably break extracellular salt bridge gating interactions involving Glu222 on TM3. Moreover, Tyr589 on TM10 participates in Cl^−^ binding at the so called S_Cl2_ site^26^, so coordination (or dissociation) of Cl^−^ to this site may also contribute to movement of TM10. Conversely, inward movement of TM8 pivoted at the extracellular end, coupled with a concerted movement of a highly conserved short helix within the ICL1, leads to occlusion of the cytoplasmic exit. Such opposite motions of TM3 and TM8 during transition among distinct transport states appear to be directly coupled as these two long tilted helices lie next to each other. In Na^+^-dependent CCC transporters, TM8 also bears two Na^+^-coordinating residues, so it is conceivable that dynamic formation and rupture of the Na^+^ site may induce movements of TM8 during ion transport as seen in our KCC1 structures.

**Figure 5.**
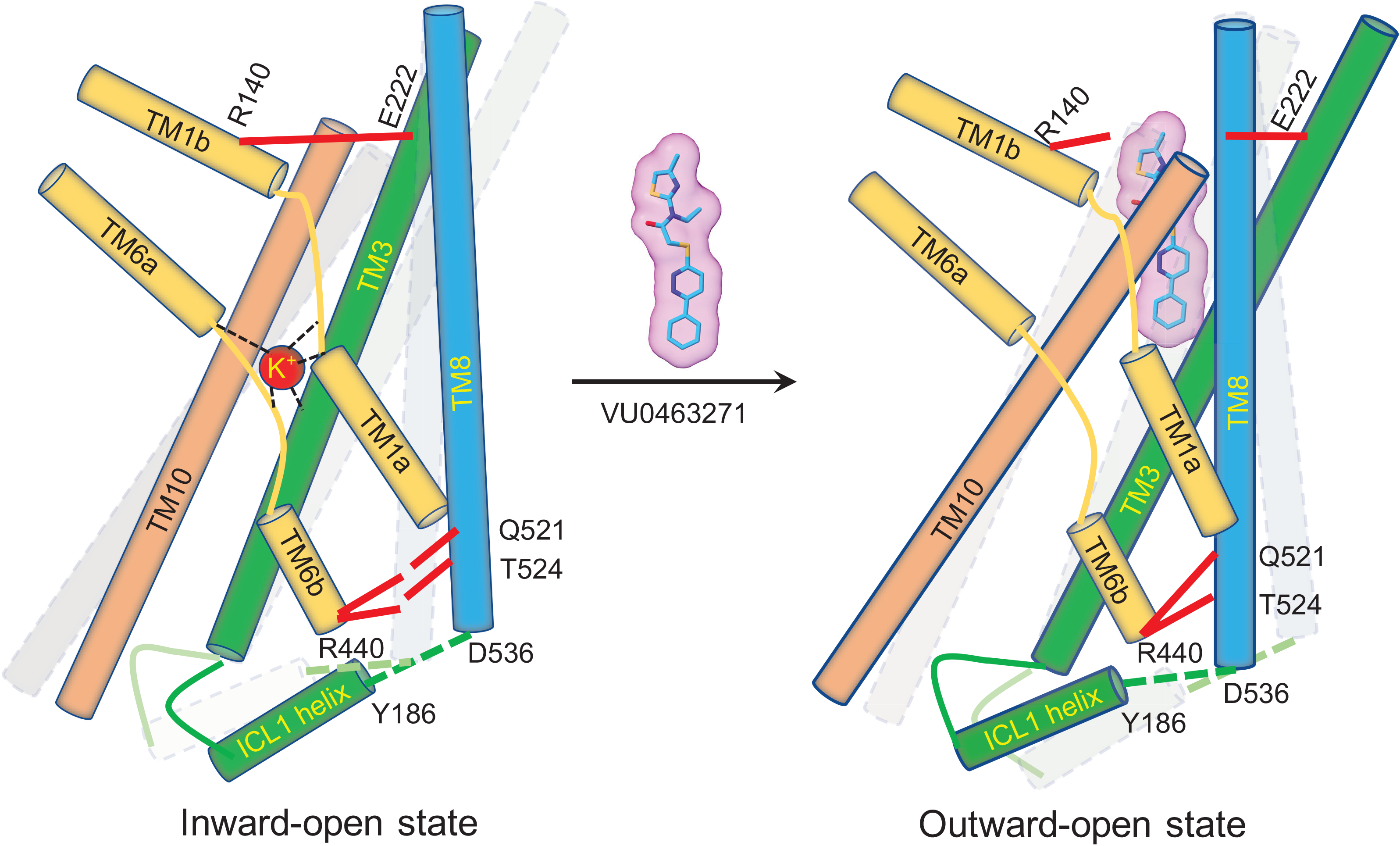
Model of gating interactions and conformational changes in KCC1. In the inward-open state, the intracellular gate is open as the gating interactions (R440-Q521 and R440-T524) are broken, while the extracellular gate is closed by the salt bridge R140-E222. The VU0463271 inhibitor arrests KCC1 in an outward-open state by wedging into the extracellular ion permeation path. VU0463271 breaks the extracellular gate, disrupts K^+^ coordination, and triggers a concerted movement of TM3, TM8, TM10, and the ICL1 helix, fostering gating interactions that close the intracellular vestibule.

Several members of the CCC family are the molecular targets of anti-hypertensive loop and thiazide diuretics and elucidating their sites and mechanisms of actions represents an important goal for further improving specificity and potency of these widely prescribed drugs via rational medicinal chemistry campaigns^28,29^. Here we report the first structure of a CCC transporter inhibited by a small molecule compound, highlighting the extracellular ion permeation path as an important molecular site for inhibition of CCC transporters. Diuretics may also trap NKCC and NCC transporters in outward-open conformations from the extracellular side as they are charged molecules and thus possibly cannot diffuse freely across the membrane barrier. We expect that our outward-open KCC1 structure will help to calculate outward-open NKCC structures based on the existing inward-open structures^24,25^. Such outward-open structures may serve as better models for computational docking campaigns to determine the sites and modes of action of various loop diuretics than the existing inward-open structures.

## Methods

### Expression and purification of human KCC1

The full-length human KCC1 isoform A gene (NG_033098.1) was amplified from human cDNA and cloned into a modified pFastBac1 vector downstream of the human cytomegalovirus (CMV) promotor. A maltose binding protein (MBP) followed by a tobacco etch virus (TEV) protease cleavage site was fused immediately before the N-terminus of KCC1. This construct was expressed in HEK293S GnTI^−/−^ cells (ATCC CRL-3022) using the BacMam system as described^46^. In brief, HEK293S GnTI^−/−^ cells, grown in suspension in Freestyle 293 expression medium (Invitrogen, Carlsbad, CA) at 37 °C in an orbital shaker, were transduced with human KCC1 baculoviruses when cell density reached ~ 2×10^6^/ml. 8 to 12 hours post transduction, sodium butyrate was added to the culture to a final concentration of 5 mM to enhance protein expression; temperature was reduced to 30 °C. Cells were harvested 72 hours post transduction and flash frozen in liquid nitrogen and stored at −80°C until use. All protein purification steps were carried out at 4°C unless stated otherwise. Membrane proteins were extracted for 1 hour at room temperature in a buffer composed of (in mM) 50 HEPES (pH7.4), 75 KCl, 75 NaCl, 0.5 tris(2-carboxyethyl)phosphine (TCEP), 3 Lauryl Maltose Neopentyl Glycol (MNG-3), and 0.6 Cholesteryl Hemisuccinate Tris Salt (CHS), 0.5 phenylmethylsulfonyl fluoride (PMSF), 5 μg/ml leupeptin, 1.4 μg/ml pepstatin A, 2 μg/ml aprotinin, and 10% glycerol. The supernatant was collected after centrifugation at 18,000 rpm for 30 minutes and then incubated with amylose resin (New England BioLabs, Ipswich, MA) for 2 hours. Human KCC1 protein was eluted from amylose resin with buffer composed of (in mM) 20 HEPES (pH7.4), 150 KCl, 0.5 TCEP, 0.5 MNG-3, 0.1 CHS, 20 maltose. MBP fusion tag was removed by incubation with TEV protease for overnight. Human KCC1 was further separated with a Superose 6 column using buffer composed of (in mM) 20 HEPES (pH7.4), 150 KCl, 0.5 TCEP, 25 × 10^−3^ MNG-3, and 5 × 10^−3^ CHS, and peaks corresponding to human KCC1 were collected and concentrated for cryo-EM analyses. For preparing human KCC1 bound with VU0463271, human KCC1 was purified with a Superose 6 column using a K^+^ free buffer composed of (in mM) 20 HEPES (pH 7.4), 150 NaCl, 0.5 TCEP, 25 × 10^−3^ MNG-3, and 5 × 10^−3^ CHS, 25 × 10^−3^ VU0463271. The peaks corresponding to KCC1 were collected and supplemented with VU0463271 to a final concentration of 75 μM to incubate for 30 minutes at room temperature prior to cryo-EM analyses.

### Proteoliposome preparation and Tl^+^ flux assay

1-palmitoyl-2-oleoyl-sn-glycero-3-phosphoethanolamine (POPE) and 1-palmitoyl-2-oleoyl-sn-glycero-3-phosphoglycerol (POPG) (Avanti Polar Lipids) were mixed at 3:1 molar ratio, dried under Argon and vacuumed for 2 hours to remove chloroform. The lipid was resuspended in 20 mM HEPES, pH7.5, 100 mM NaCl to a final concentration of 10 mg/ml, sonicated to transparency and incubated with 40 mM n-decyl-β-D-maltoside (DM, Anatrace) for 2 hours at room temperature under gentle agitation. Wild type or mutant human KCC1 was added at 1:100 by weight protein to lipid ratio. The detergent was removed by dialysis at 4 °C against the inside buffer (20 mM HEPES, pH7.5, 100 mM Na-Gluconate) in 20 kDa molecular weight cutoff dialysis cassettes. Buffer was changed every 24 hours, and after 4 days of dialysis, the proteoliposomes were harvested, aliquoted, and frozen at −80 °C.

A Tl^+^-sensitive fluorescence dye, ThaLux-AM (WaveFront Biosciences), was converted to its membrane impermeable form following a published protocol^47^. After the conversion, the pH was adjusted to 7.5 with H_2_SO_4_. The working concentration of the dye was 50 μM. The dye was incorporated into proteoliposomes by three cycles of freeze-thaw, and then extrusion through a 400 nm filter (NanoSizer™ Extruder, T&T Scientific Corporation). Excess dye was removed through a desalting column (PD-10, GE Healthcare) equilibrated with the outside buffer (20 mM HEPES, pH7.5, 96 mM Na-Gluconate, 4 mM NaCl).

Proteoliposomes were added to a quartz cuvette, and fluorescence was monitored at 520 nm with an excitation wavelength of 494 nm (FluoroMax-4, HORIBA). The transport was initiated by addition of 0.5 mM Tl_2_SO_4_. When testing an inhibitor, desired concentration of the inhibitor was added to the cuvette and incubated for at least 5 minutes before initiation of the transport.

The rate of ion transport was estimated by the slope of a fluorescence trace in the first 30 seconds. Percent of activities were calculated by defining 0% for liposomes with no protein and 100% for liposomes with human KCC1 and in the absence of an inhibitor and plotted versus inhibitor concentrations. The median inhibition concentration (IC_50_) was obtained by fitting the data to Y=1/(1+10^(X-LogIC_50_)) in the GraphPad Prism 8.0 software.

### Cryo-EM data acquisition

2.5 μl of KCC1 sample at ~ 6 - 8 mg/ml, with or without the VU0463271 inhibitor, was applied to glow-discharged Quantifoil 2.0/2.0 holey, 200 mesh carbon grids. Grids were plunge frozen in liquid ethane using a Vitrobot Mark III (FEI) set to 4°C, 80% relative humidity, 20 s wait time, −1 mm offset, and 2.5 s blotting time. Data were collected on a Krios (FEI) operating at 300 kV equipped with the Gatan K3 direct electron detector at the University of Utah and Pacific Northwest Cryo-EM Center (PNCC). Images were recorded using SerialEM^48^, with a defocus range between −1.0 to −3.5 μm. For the KCC1 bound with VU0463271 sample, we recorded movies in super-resolution counting mode with a physical pixel size of 1.035 Å, at a dose rate of 0.75 e^−^/Å^2^/frame with a total exposure of 67 frames, giving a total dose of 50 e^−^/Å^2^. For the KCC1 in 150 mM KCl sample, we also recorded movies in super-resolution counting mode with a physical pixel size of 1.092 Å, at a dose rate of 1 e^−^/Å^2^/frame with a total exposure of 40 frames, giving a total dose of 40 e^−^/Å^2^.

### Image processing 3D reconstruction and model building

Movie frames were aligned, dose weighted, and then summed into a single micrograph using MotionCor2^49^. CTF parameters for micrographs were determined using the program CTFFIND4^50^. Approximately 4000 particles were manually boxed out in cryoSPARC to generate initial 2D averages, which were then used as template to automatically pick particles from all micrographs in cryoSPARC^51^. For the KCC1/VU0463271 dataset, a total of 2,514,195 particles were extracted and then subjected to one round of 2D classification in cryoSPARC. ‘Junk’ particles that were sorted into incoherent or poorly resolved classes were rejected from downstream analyses. The remaining 446,018 particles from well resolved 2D classes were pooled and the subjected to another round of 2D classification in cryoSPARC, and the resulting 242,520 good particles were used to calculate a *de novo* model in cryoSPARC without imposing any symmetry. At this point, the 446,018 particles from the first round of 2D classification were exported into RELION^52^ for 3D classification with C2 symmetry imposed and using the cryoSPARC *de novo* map as the starting model. The 99,223 particles from one good 3D class were then subjected to non-uniform refinement in cryosparc and yielded a 4.1 Å map. At this point, the 99,223 particles were again exported into RELION and the orientation parameters calculated in cryoSPARC were used for CTF refinement and Bayesian polishing^53^. The polished 99,223 particles were again imported back to cryoSPARC for non-uniform refinement that calculated a final map of 3.63 Å resolution when solvent noise and detergent micelle belt were masked out.

For the KCC1 in 150 mM KCl sample, a total of 4,073,077 particles were extracted and subjected to one round of 2D classification in cryoSPARC, resulting in 1,151,775 good particles. Subsequently, one round of heterogenous 3D classification was performed in cryoSPARC without imposing any symmetry using a fly KCC map (unpublished) filtered to 20 Å map as an initial model. A single class showing clear secondary structural features and best map connectivity was selected and the 306,308 particles in this class were subjected to non-uniform refinement in cryoSPARC and yielded a 3.7 Å map. At this point, the 306,308 particles were exported into RELION for 3D classification without alignment (i.e., the Euler angles and in plane shifts calculated by non-uniform refinement in cryoSPARC were maintained) or with local search. In both 3D classification, the extracellular domains of KCC1 were masked out. The good 3D classes from these two 3D classification calculations were combined and duplicated particles were removed. The resulting 79,153 particles were subjected to CTF refinement and Bayesian polishing in RELION and then imported back to cryoSPARC for non-uniform refinement that calculated a final map of 3.25 Å resolution when solvent noise and detergent micelle belt were masked out.

The maps was locally sharpened in cryoSPARC with an overall b factor of ~ −120 Å^2^ (KCC1 in 150 mM KCl) or −175 Å^2^ (KCC1 bound with UV0463271) for model building in Coot^54^. The human KCC1 transmembrane structure (PDB: 6KKR) was dock into both maps and adjusted in Coot. VU0463271 was docked into the KCC1 map via Autodock Vina and the final pose was further adjusted manually according to the local chemical environment^55^. The cytoplasmic C-terminal domain was built *de novo* into the KCC1 map determined in the presence of 150 mM KCl. Most of the KCC1 C-terminal domain can be unambiguously modeled, but the region encompassing residues 882-929 was built tentatively due to poorly resolved densities. The model was refined in real space using PHENIX^56^, and assessed in Molprobity (Supplementary Table 1)^57^. FSC curves were then calculated between the refined model *versus* summed half maps generated in cryoSPARC, and resolution was reported as FSC=0.5 (Extended Data Figure1 and S3). UCSF Chimera was used to visualize and segment density maps, and figures were generated using Chimera and Pymol. HOLE program was used to calculate solvent accessible pathways^58^.

## Data and software availability

The cryo-EM maps of human KCC1 have been deposited in the Electron Microscopy Data Bank (EMDB) with the accession codes EMD-***** (VU0463271 bound) and EMD-***** (150 mM KCl). The atomic coordinates for the corresponding maps have been deposited in the Protein Data Bank (PDB) with the accession codes ****(VU0463271 bound) and **** (150 mM KCl). All other data and reagents that support the findings of this study are available from the corresponding author upon request.

## Author contributions

Conceptualization, Y.Z. and E.C. designed cryo-EM studies, and J.S. and M.Z. designed functional analyses; Investigation, Y.Z. carried out cryo-EM experiments, J.S. performed transport assays in liposomes, and Q.W. contributed to model building; Writing, E.C., Y.Z., Q.W., J.S., and M.Z‥

## Declaration of interest

The authors declare no competing interest.

## Acknowledgements

This work was primarily supported by the start-up fund from the University of Utah. E.C. is a Pew Scholar supported by the Pew Charitable Foundation. We thank Anita Orendt, Irvin Allen, Martin Cuma, and other staff members at the Utah Center for High Performance Computing for computational support. We are grateful to David Timm and David Belnap for data collection at the Electron Microscope Core at the University of Utah. The Electron Microscope Core at the University of Utah was supported by a grant from the Beckman Foundation. A portion of this research was supported by NIH grant U24GM129547 and performed at the PNCC at OHSU and accessed through EMSL (grid.436923.9), a DOE Office of Science User Facility sponsored by the Office of Biological and Environmental Research. We thank Claudia Lopez, Harry Scott, Janette Myers, Drew Gingerich, and other staff members at the PNCC for data collection and technical support.

---

**Extended Data Figure 1.**
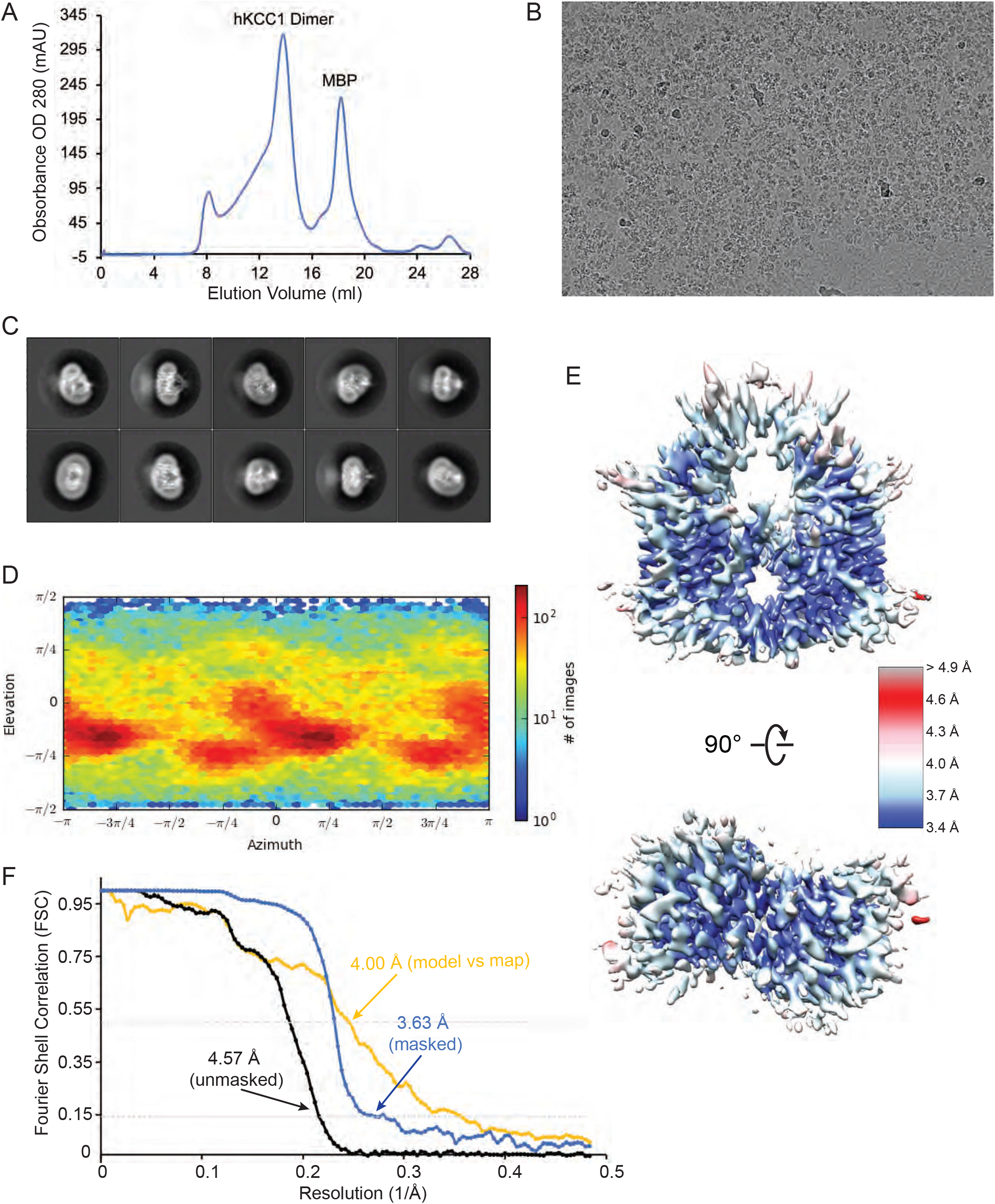
Cryo-EM structure of human KCC1 bound with VU0463271. (a) Size exclusion chromatogram of the full-length human KCC1. (b) A representative micrograph of KCC1/VU0463271 recorded with a Titan Krios microscope. (c) 2D class averages of KCC1/VU0463271 show well-resolved structural features for the transmembrane domain, but not for the cytosolic domains. (d) Angular distribution plot of all particle projections as outputted by cryoSPARC. (e) Local resolutions calculated in cryoSPARC. (f) Gold-standard FSC curves calculated after cryoSPARC non-uniform refinement.

**Extended Data Figure 2.**
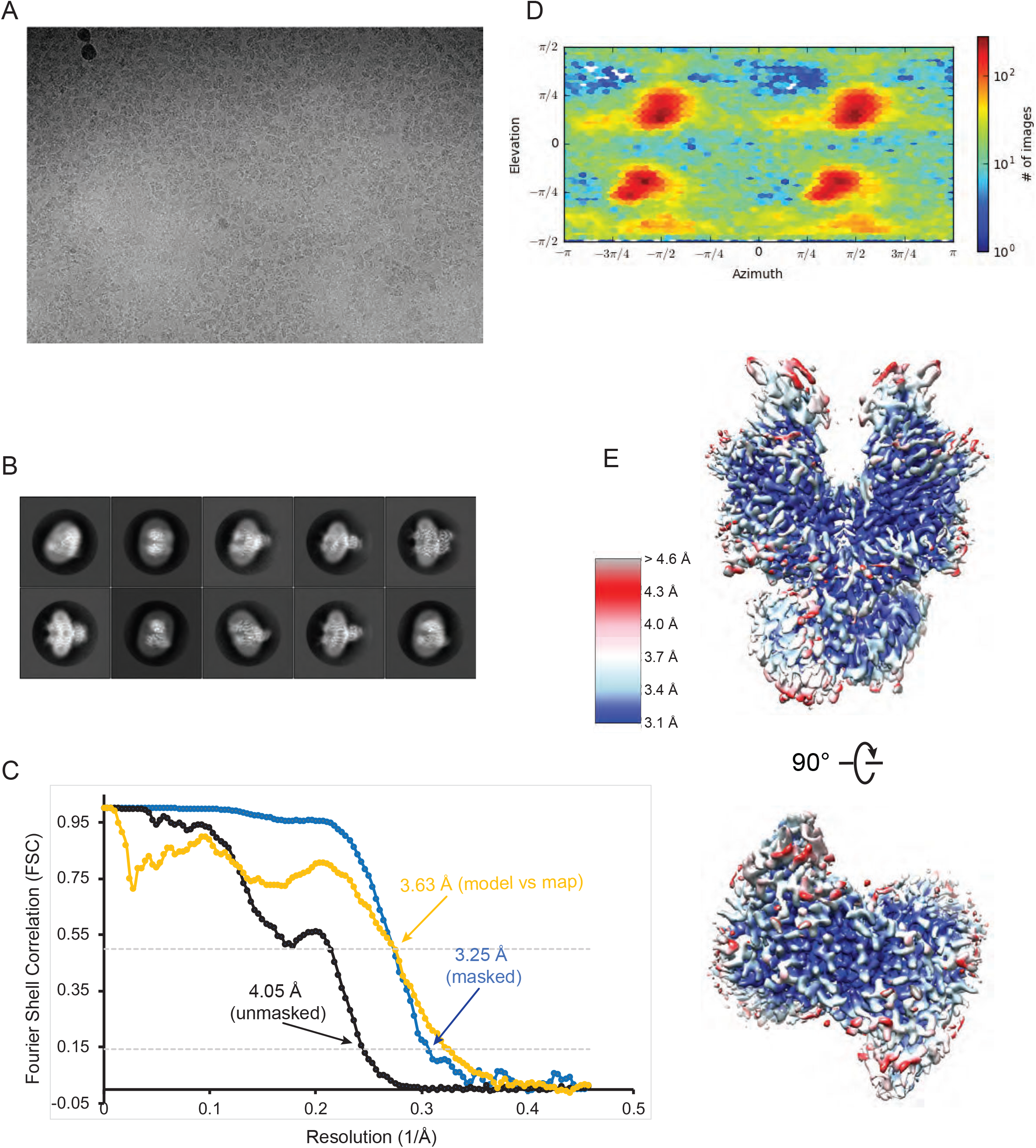
Cryo-EM structure of human KCC1 in 150 mM KCl. (a) A representative micrograph of KCC1 in 150 mM KCl recorded with a Titan Krios microscope. (b) 2D class averages of KCC1 in 150 mM KCl show well-resolved structural features. (c) Gold-standard FSC curves calculated after cryoSPARC non-uniform refinement. (d) Angular distribution plot of all particle projections as outputted by cryoSPARC. (e) Local resolutions calculated in cryoSPARC.

**Extended Data Figure 3.**
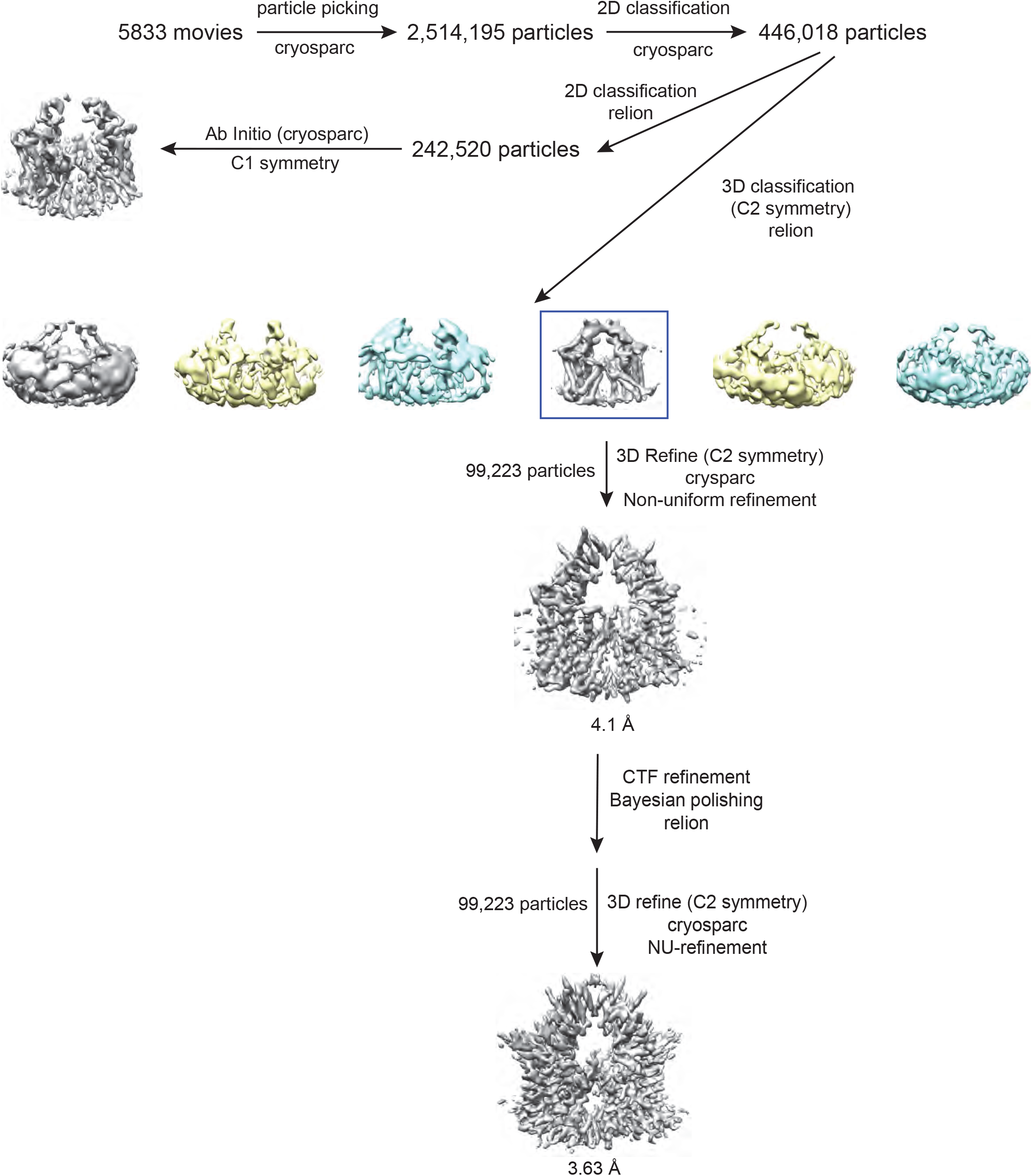
3D Reconstruction of human KCC1 bound with VU0463271. Flow chart of image processing for the human KCC1 bound with VU0463271 dataset.

**Extended Data Figure 4.**
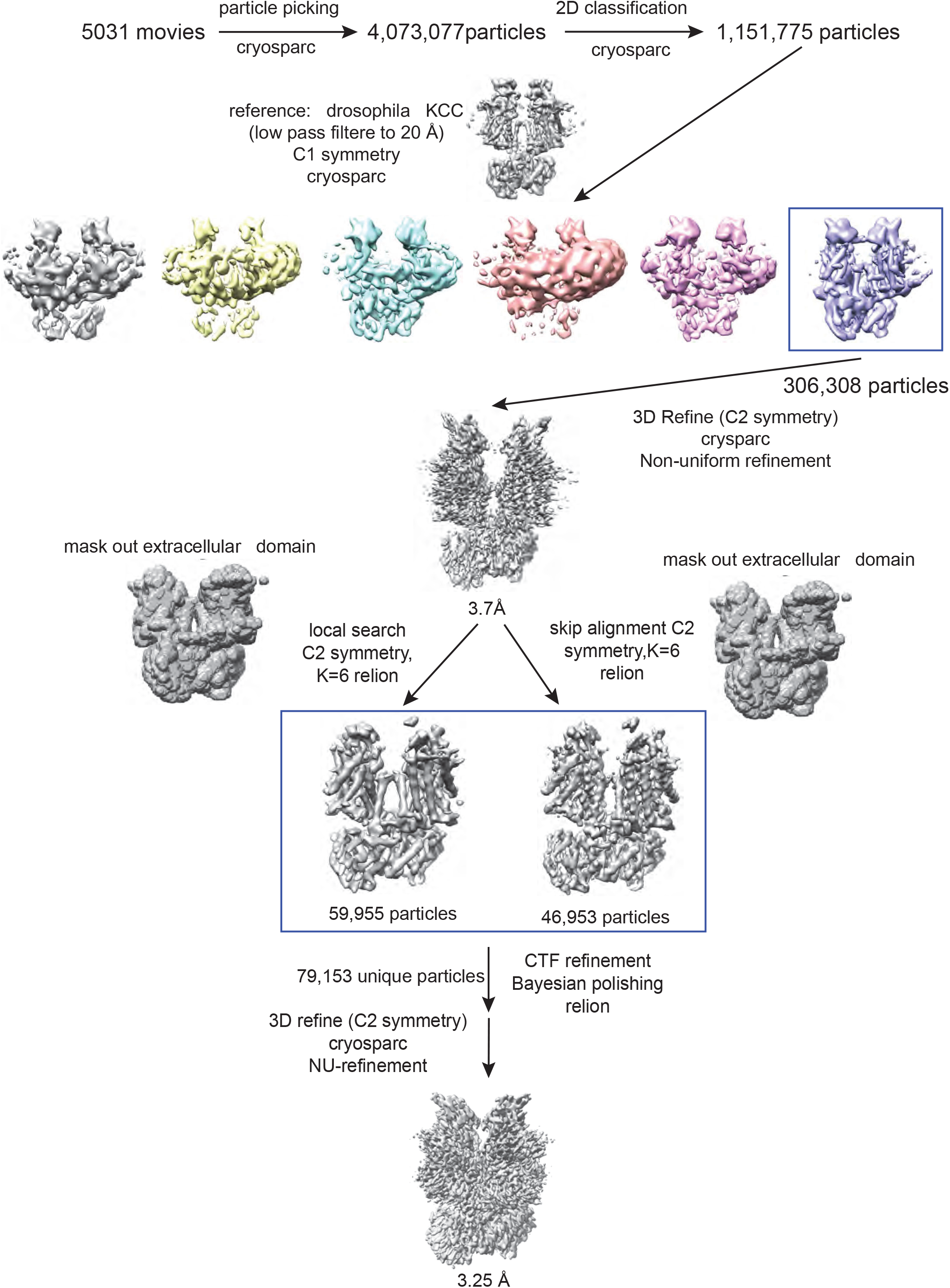
3D Reconstruction of human KCC1 in 150 mM KCl. Flow chart of image processing for the human KCC1 in 150 mM KCl dataset.

**Extended Data Figure 5.**
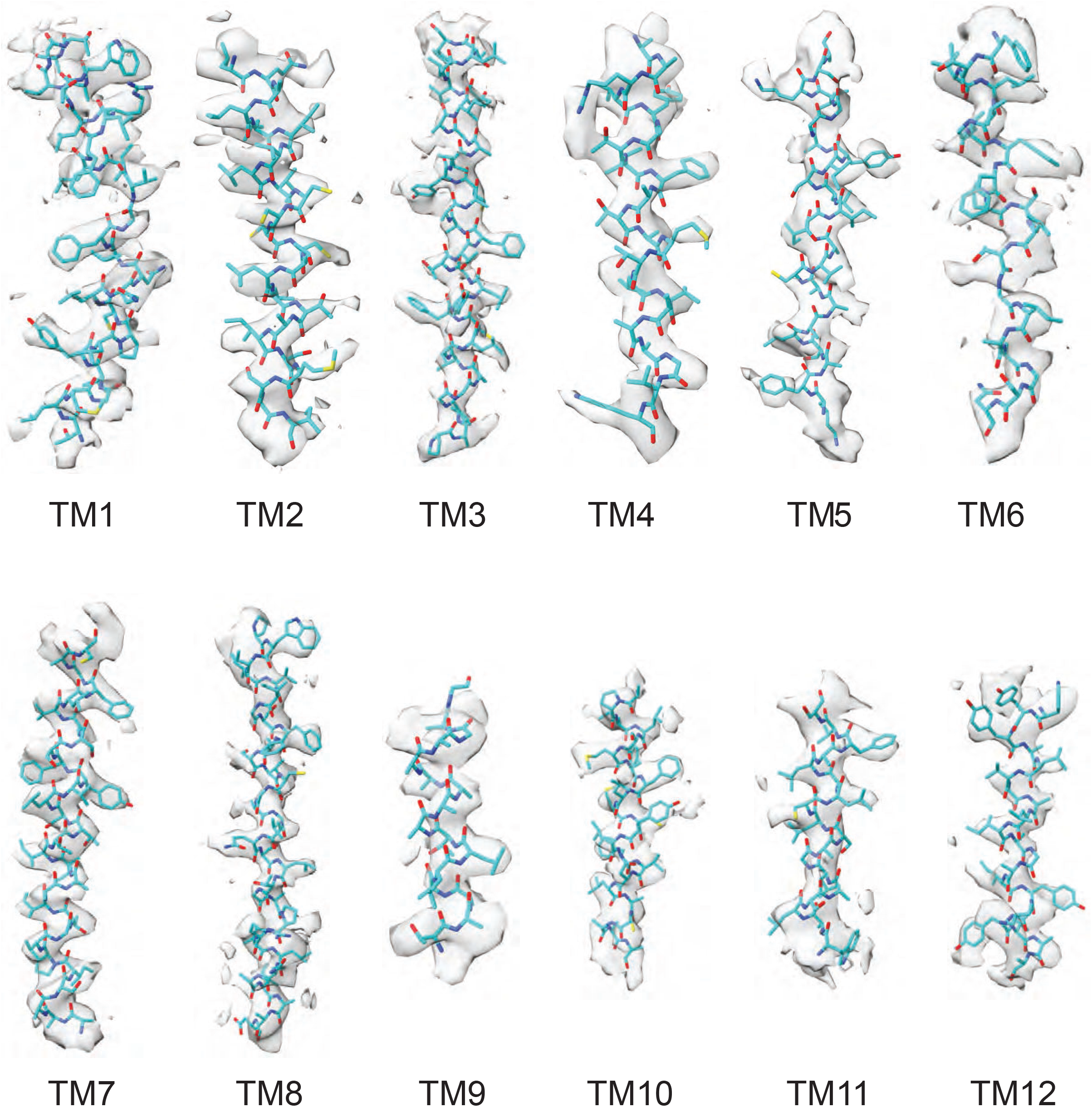
EM density maps of transmembrane helices of human KCC1 bound with VU0463271. The map is locally sharpened with an overall b factor of −175 Å^2^ in cryoSPARC. The final model shown in sticks is docked into experimental densities.

**Extended Data Figure 6.**
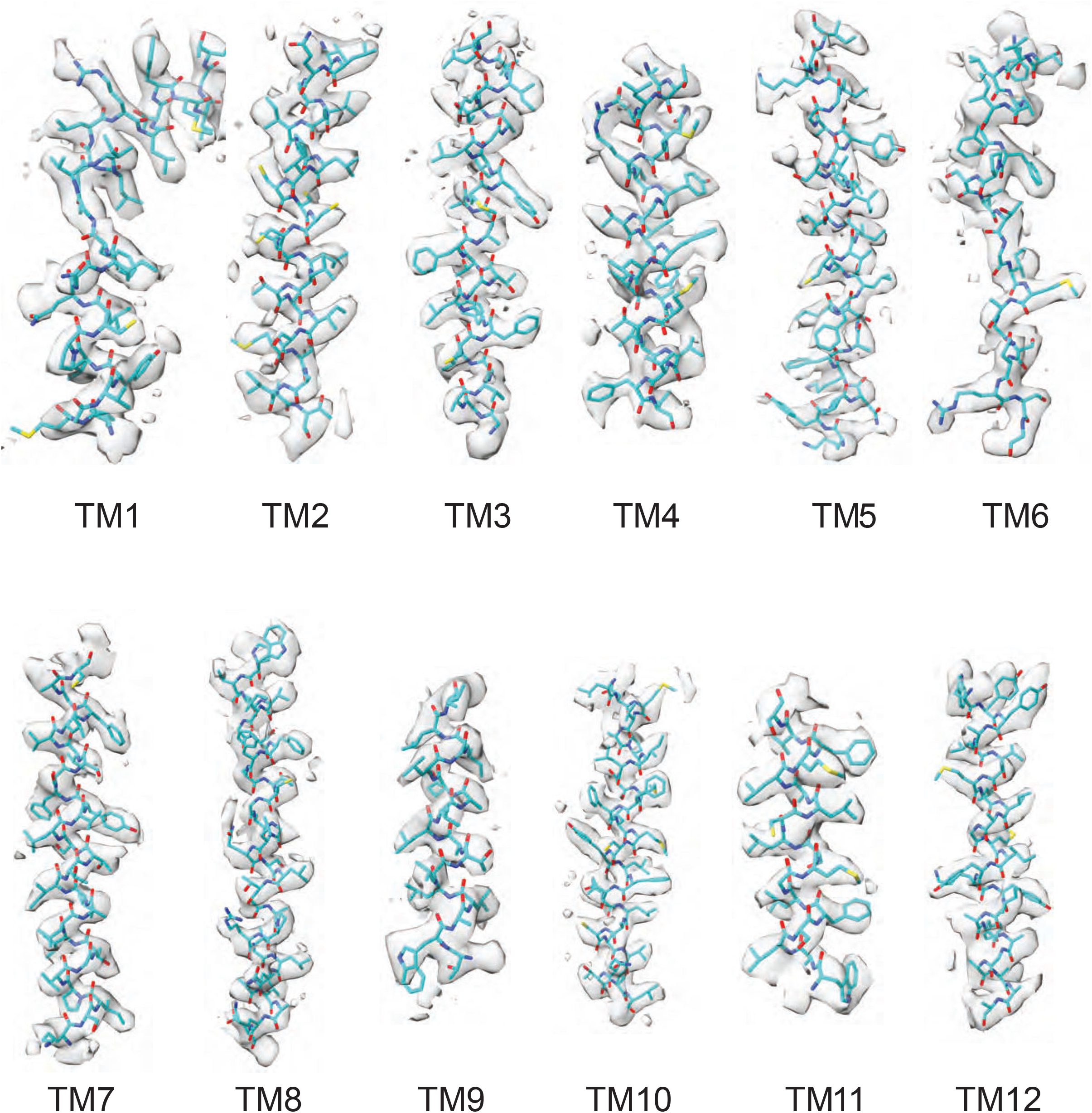
EM density maps of transmembrane helices of human KCC1 in 150 mM KCl. The map is locally sharpened with an overall b factor of −120 Å^2^ in cryoSPARC. The final model shown in sticks is docked into experimental densities.

**Extended Data Figure 7.**
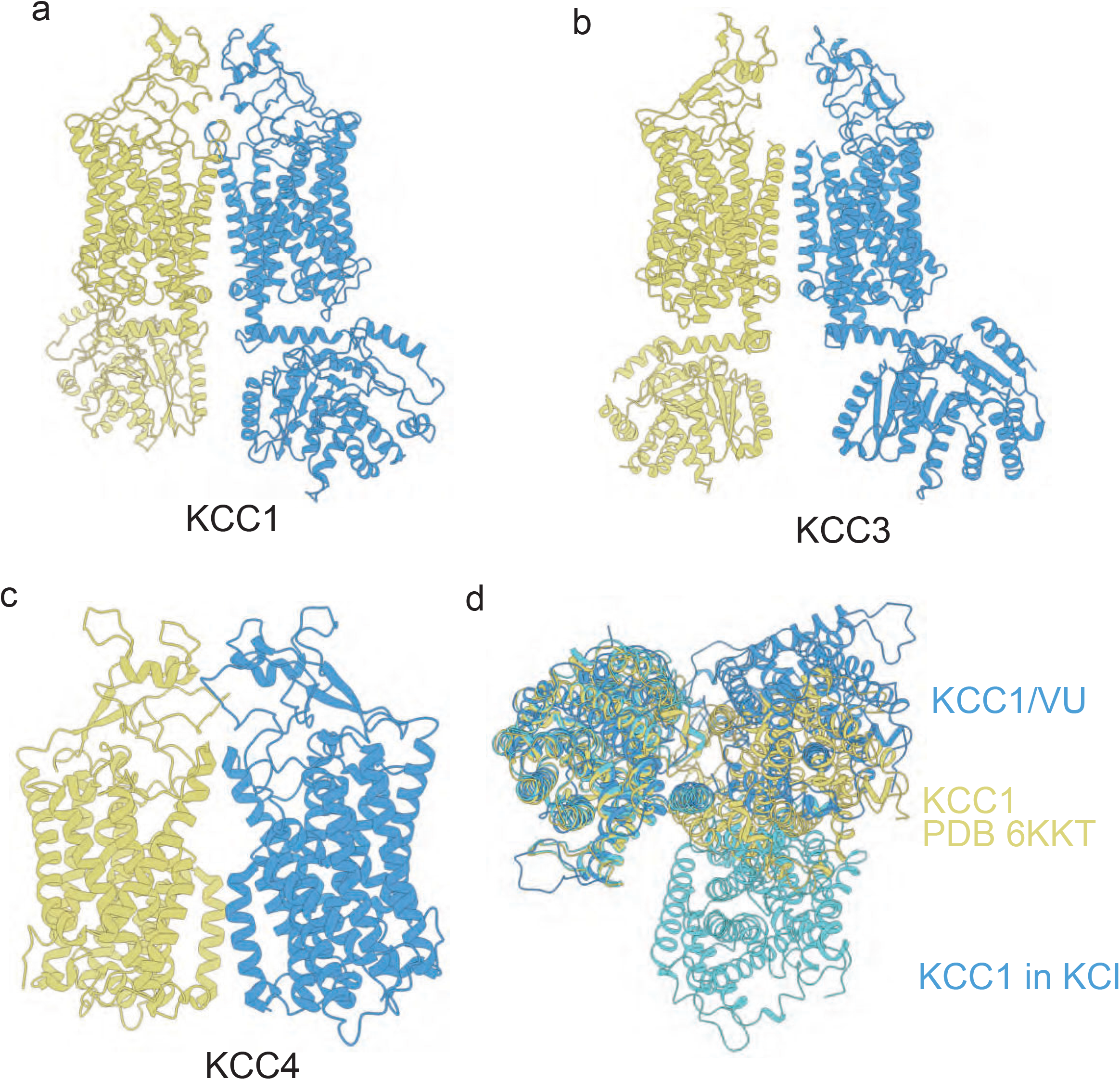
KCCs may adopt different dimeric architectures. (a) In the KCC1/VU0463271 structure, the cytosolic C-terminal domains would be separated from each other if each KCC1 subunit retains the architecture as seen in KCC1 in the presence of 150 mM KCl. (b-c) Hypothetic KCC3 and KCC4 dimers modeled with the KCC1 architecture in the presence of VU0463271. Their large extracellular domains may similarly participate in homotypic interactions as seen in the KCC1/VU0463271 structure. (d) Three human KCC1 structures, two of which are determined in this study, are aligned based on one subunit, highlighting distinct dimeric organizations as the second subunit is differently oriented in relation to the first subunit.

**Extended Data Figure 8.**
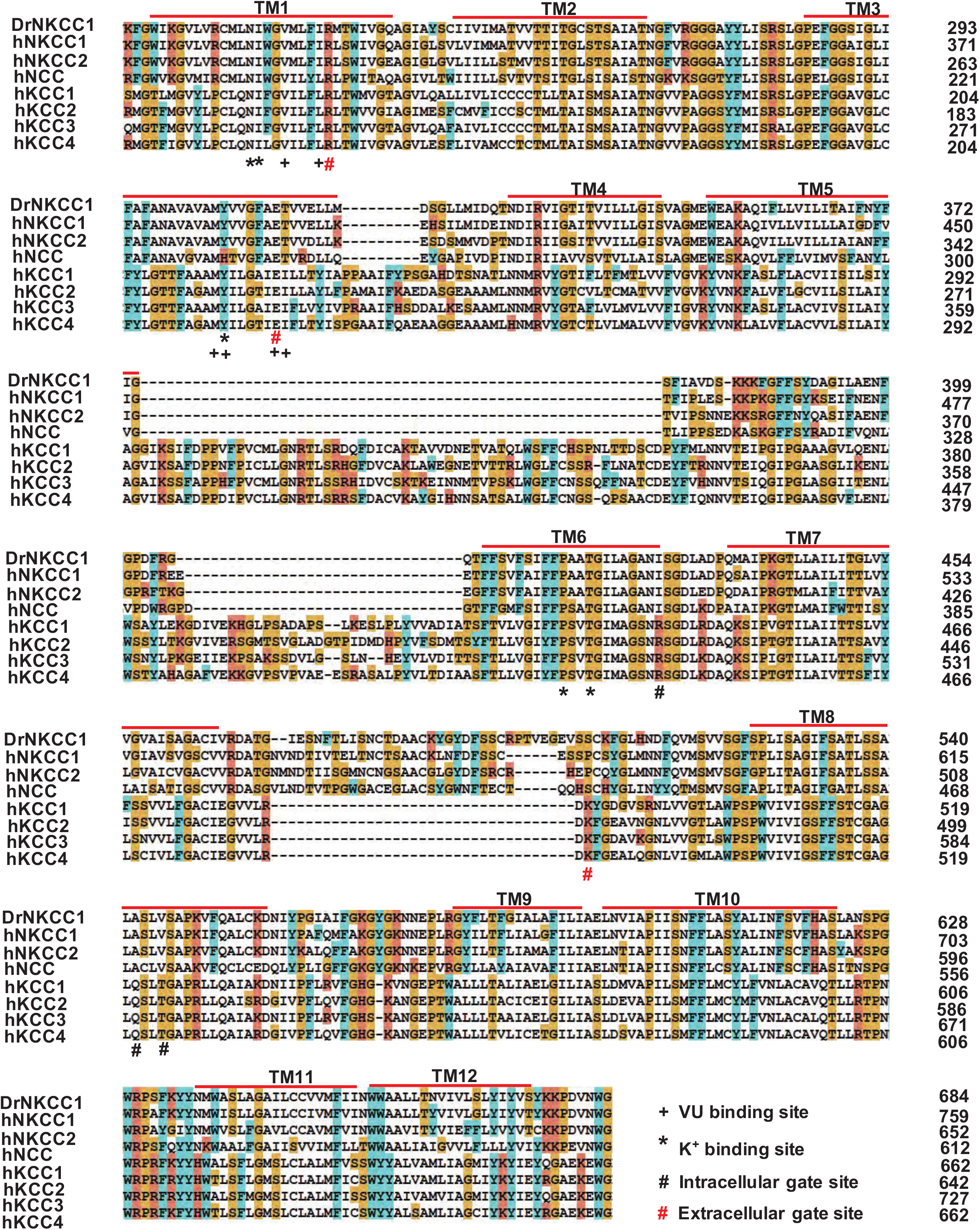
Alignment of sequences of CCC transporters. Functional important residues are highlighted as indicated.

**Extended Data Figure 9.**
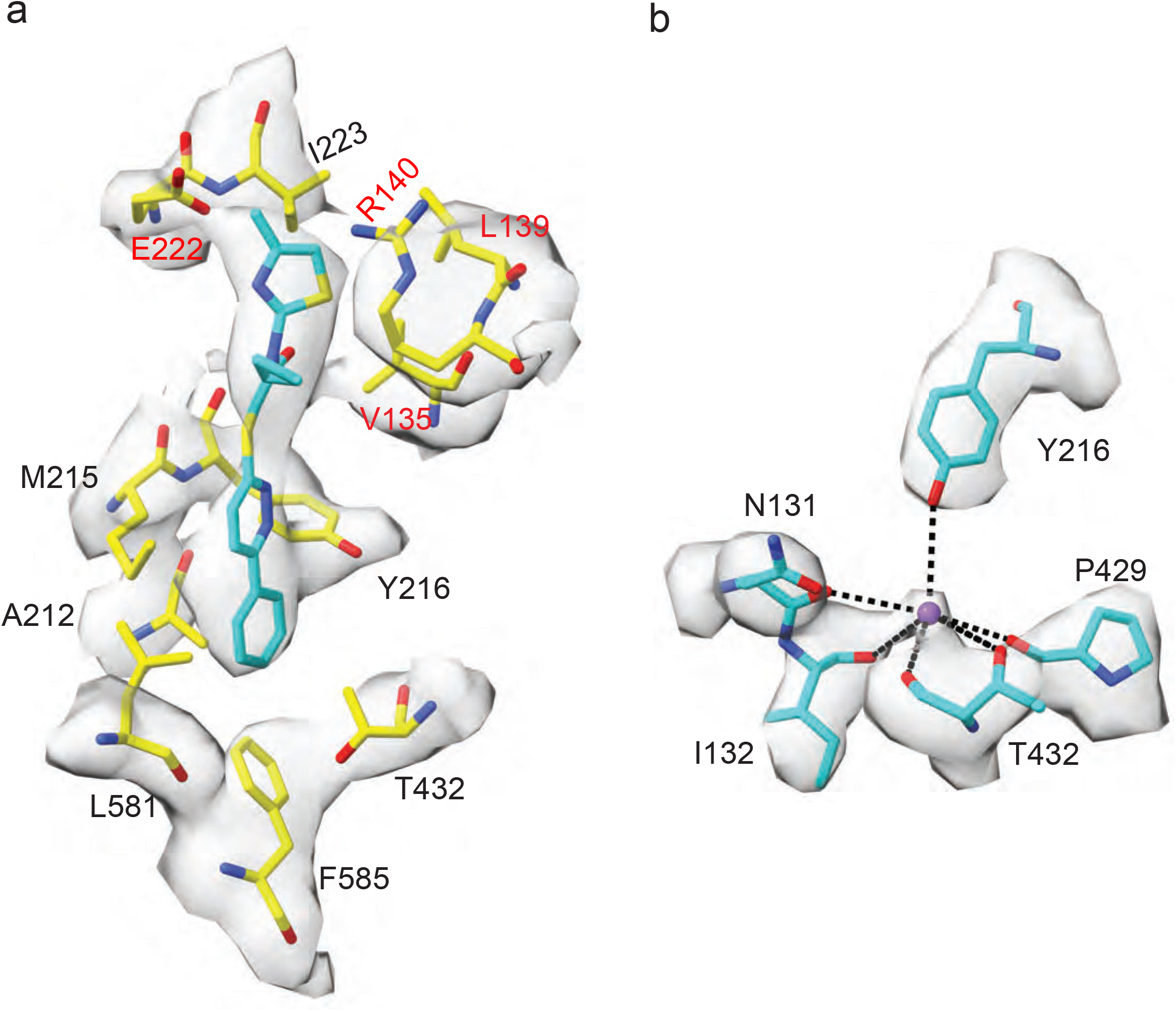
Binding sites for VU0463271 and K^+^ overlap in human KCC1. VU0463271 and K^+^ are shown as stick and purple sphere, respectively.

**Extended Data Table 1.** Statistics of cryo-EM data acquisition and processing and model building.

| **Data collection/Processing** |  |  |
| --- | --- | --- |
|  | Human KCC1 bound with VU0463271 | Human KCC1 in a KCl containing buffer |
| Microscope | Titan Krios/K3 camera | Titan Krios/K3 camera |
| Voltage (kV) | 300 | 300 |
| Defocus range ($\mu\text{m}$) | -0.7 – -3.5 | -0.7 – -2.9 |
| Pixel size ($\text{\AA}$) | 1.035 | 1.092 |
| Total electron dose ($\text{e}^-/\text{\AA}^2$) | 50 | 40 |
| Exposure time (s) | 8 | 2.4 |
| Number of movies | 5,833 | 5,031 |
| Number of frames per movie | 67 | 40 |
| Initial particle number | 2,514,195 | 4,073,077 |
| Final particle number | 99,223 | 79,153 |
| Resolution (unmasked, $\text{\AA}$) | 4.57 | 4.05 |
| Resolution (masked, $\text{\AA}$) | 3.63 | 3.25 |
| **Refinement and Validation** |  |  |
| Number of atoms | 7,260 | 12,744 |
| R.M.S deviation |  |  |
| Bond length ($\text{\AA}$) | 0.006 | 0.006 |
| Bond angles ($^\circ$) | 1.023 | 0.985 |
| Ramachandran |  |  |
| Favored (%) | 91.73% | 92.94% |
| Allowed (%) | 8.27% | 7.06% |
| Outlier (%) | 0.00% | 0.00% |
| Molprobity score | 2.86 | 2.38 |

## Notes

### Competing Interest Statement

The authors have declared no competing interest.

